# Intermembrane space-localized twin Cx_9_C motif-containing TbTim15 is an essential subunit of the single mitochondrial inner membrane protein translocase of trypanosomes

**DOI:** 10.1101/2024.01.31.578135

**Authors:** Corinne von Känel, Silke Oeljeklaus, Christoph Wenger, Philip Stettler, Anke Harsman, Bettina Warscheid, André Schneider

## Abstract

All mitochondria import >95% of their proteins from the cytosol. This process is mediated by protein translocases in the mitochondrial membranes, whose subunits are generally highly conserved. Most eukaryotes have two inner membrane protein translocases (TIMs) that are specialized to import either presequence-containing or mitochondrial carrier proteins. In contrast, the parasitic protozoan *Trypanosoma brucei* has a single TIM complex consisting of one conserved and five unique subunits. Here, we show that the trypanosomal TIM complex contains an additional trypanosomatid-specific subunit, designated TbTim15. TbTim15 is associated with the TIM complex, lacks transmembrane domains and localizes to the intermembrane space. TbTim15 is essential for procyclic and bloodstream forms of trypanosomes. It contains two twin CX_9_C motifs and mediates import of both, presequence-containing and mitochondrial carrier proteins. While the precise function of TbTim15 in mitochondrial protein import is unknown, our results are consistent with the notion that it may function as an import receptor for the non-canonical trypanosomal TIM complex.

## Introduction

Mitochondria or mitochondria-derived organelles are hallmarks of eukaryotes and fulfil various essential functions^1^. Due to their endosymbiotic origin, mitochondria contain their own genome, which however, only encodes for a small fraction of mitochondrial proteins^2^. Thus, >95% of all mitochondrial proteins are encoded in the nucleus, synthesized in the cytosol, and subsequently imported into mitochondria^3,4^. Most experimental studies on mitochondrial protein import have been done in the yeast *Saccharomyces cerevisiae* and in mammals, both of which belong to the eukaryotic supergroup Opisthokonta^5^. These studies have shown that the majority of proteins enter mitochondria through a common entry gate, the translocase of the outer mitochondrial membrane (TOM)^6,7^. Depending on distinct targeting signals, the imported substrates are then sorted to their submitochondrial destination. Yeast and most other eukaryotes have two translocases of the inner mitochondrial membrane (TIM), the TIM22 and TIM23 complexes^8,9^. The TIM22 complex, also referred to as the carrier translocase, mediates the insertion of proteins into the inner mitochondrial membrane (IM) which have multi-spanning transmembrane domains (TMDs), such as mitochondrial carrier proteins (MCPs)^10,11^. The TIM23 complex, also termed presequence translocase, imports presequence-containing precursor proteins across or into the IM^12–14^, which represent 60-70% of all mitochondrial proteins^15^. To that end, TIM23 associates with the matrix-exposed presequence translocase-associated motor (PAM). The yeast PAM consists of five essential components^13,16^: the mitochondrial heat shock protein 70 (mHsp70)^17,18^, the J-domain containing co-chaperones Pam18^19,20^ and Pam16^21^, Tim44 which tethers mHsp70 to TIM23^22^ and the nucleotide exchange factor Mge1^23–25^.

During the last decade, the unicellular parasite *Trypanosoma brucei,* a member of the eukaryotic supergroup of the Discoba^5^, has emerged as an experimentally highly accessible, non-opisthokont model system for mitochondrial biology. Studies on mitochondrial protein import in *T. brucei* have shown that the trypanosomal import systems are highly diverged compared to other eukaryotes^26–28^. Accordingly, the trypanosomal TOM complex, referred to as atypical TOM (ATOM), shares only two of its seven subunits with other eukaryotes^29–31^. However, the largest differences to other eukaryotes concern the IM translocases. Epitope-tagged import substrates, stalled within either the presequence translocase or the carrier translocase, were used to show that *T. brucei* has a single TIM complex only. This single TIM, with minor compositional variations, can import presequence-containing proteins as well as MCPs^32^. It consists of the essential integral IM proteins TbTim17, TbTim62, TbTim42 and a medium chain length acyl-CoA dehydrogenase (ACAD)^32–35^. TbTim17 is an orthologue of Tim22, the pore-forming subunit of the TIM22 complex in yeast^8,9,36^. It is the only integral membrane TIM subunit that shares homology with components of the IM translocase of other eukaryotes^32,34^. Furthermore, the single TIM complex is associated with six intermembrane space (IMS)-localized small Tim chaperones, termed Tim8-13, Tim9, Tim10, TbTim11, TbTim12 and TbTim13^32,37,38^. The trypanosomal small Tims show homology to small Tim chaperones of other eukaryotes that specifically interact with the TIM22 complex and escort hydrophobic substrate across the aqueous IMS. Two additional integral membrane proteins, the proteolytically inactive rhomboid-like proteins TimRhom I and TimRhom II, are associated with the trypanosomal TIM complex. Interestingly, they appear to be selectively required for import of presequence-containing proteins^32^. Finally, import of presequence-containing proteins requires the trypanosomal TIM complex to associate with an unconventional PAM module^39^. The trypanosomal PAM contains an orthologue of mHsp70 (TbmHsp70) and the kinetoplastid-specific J-domain protein TbPam27^39^ but lacks homologues of yeast Pam18, Pam16 and Tim44.

Using tagged TbPam27 as the bait in a quantitative co-immunoprecipitation (CoIP) experiment, we discovered two new subunits of the trypanosomal TIM complex, which were termed TbTim15 and TbTim20. Both proteins were also enriched in pull down experiments of three other TIM subunits^32,39^. TbTim15 is an IMS-localized TIM complex-associated protein that mediates import of presequence-containing and of MCPs. In contrast, TbTim20 appears to have a non-essential function and selectively associates with the active and inactive presequence translocase form of the TIM complex.

## Results

### Identification of novel trypanosomal TIM or PAM subunit candidates

TbPam27, aside from TbmHsp70, is the first experimentally analysed subunit of the trypanosomal PAM. It is closely associated with the single TIM complex, when it translocates prese-quence-containing proteins^32,39^. To identify new putative TIM or PAM subunits, we used C-terminally myc-tagged TbPam27 (TbPam27-myc) as the bait in a CoIP experiment employing stable isotope labelling by amino acids in cell culture (SILAC). Subsequently, the eluted proteins were analysed by quantitative mass spectrometry (MS) in triplicate experiments to determine protein abundance ratios.

The resulting TbPam27-myc SILAC CoIP eluate contained 74 mitochondrial proteins^40^ that were enriched more than fivefold (*Figure 1A, Table S1*). Importantly, all six essential integral membrane components of the presequence-translocase form of the TIM complex^32^ were among the top ten enriched proteins (all more than ninefold) following the bait. Furthermore, the six trypanosomal small Tim chaperones^32,37,38^ were all more than sixfold enriched (*Fig. 1A, Table S1*). To identify new putative TIM or PAM complex subunits, we generated an edgeweighted protein-protein interaction network using Cytoscape (version 3.9)^41^. For this analysis, we considered the TbPam27 SILAC CoIP and the previously published SILAC CoIPs of the TIM core subunits (TbTim17^32^, TbTim42^32^, ACAD^39^) as well as the small Tim protein TbTim13^32^. The network contains proteins that were enriched more than threefold in at least two of these SILAC CoIP experiments (*Fig. 1B*). Of the resulting cluster, we chose five proteins, Tb927.2.4445, Tb927.11.1620, Tb927.11.1010, Tb927.11.16750 and Tb927.8.3450, that have been especially highly enriched (more than fivefold) in the TbPam27 SILAC CoIP (*Fig. 1AB, Table S1*), for further experimental analysis. ^32,39^. We named Tb927.2.4445 and Tb927.11.1620, TbTim15 and TbTim20, respectively, according to their predicted molecular weights. The remaining candidates will be referred to as 1010, 16750 and 3450.

**Figure 1.**
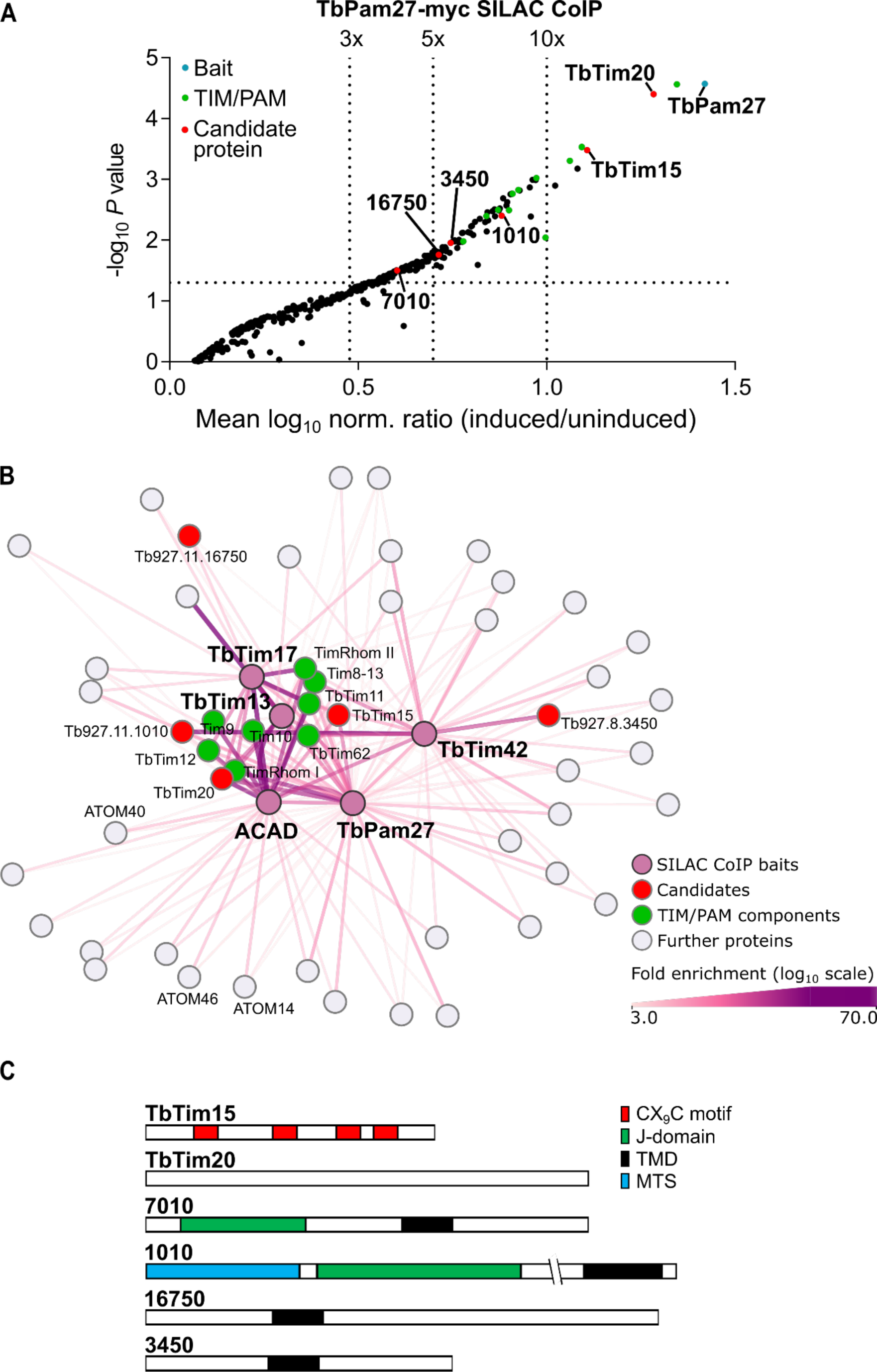
-TbPam27-myc SILAC CoIP reveals new TIM and PAM subunit candidates. **(A)** Volcano plot depicting mitochondrial proteins detected in SILAC-based quantitative mass spectrometry analysis of TbPam27-myc CoIPs. Y-axis depicts adjusted p-value (-log_10_). Differentially labelled uninduced and in-duced cells were mixed and subjected to CoIP. The vertical dotted lines in the volcano plot indicate the specified enrichment factors. The horizontal dotted line indicates an adjusted p-value of 0.05. The bait TbPam27 is highlighted in blue, TIM and PAM components in green and candidate proteins chosen for experimental analysis in red. **(B)** Edge-weighted protein-protein interaction network generated using Cytoscape (version 3.9). Network depicts proteins enriched more than threefold in at least two of the SILAC CoIPs using TbPam27, TbTim17, TbTim42, ACAD and TbTim13 as the baits (baits shown in pink). Candidates chosen for experimental analysis are shown in red, TIM and Pam subunits in green and further proteins in gray. The fold enrichment is indicated by the line-with and color intensity of the connecting lines. Candidate 7010 was enriched in the TbPam27 SILAC CoIP, but not in the other ana-lyzed TIM subunit CoIPs. Therefore, it is not part of the protein interaction network. **(C)** Schematic to scale representation of candidate proteins. *In silico* predicted domains are indicated.

Of the five candidates, only TbTim15 and TbTim20 were enriched in a SILAC pull down experiment of a tagged presequence-containing substrate that is stuck in the TIM complex (*Fig. S1A*)^32^. Whereas none of them, with the possible exception of TbTim15, which was not detected in the SILAC pull down, was recovered with a truncated tagged MCP that is stalled in the TIM complex^32^. To find out whether TbTim15 is part of the active carrier translocase, we repeated this experiment (*Fig. S1B*)^32^. The result showed that TbTim15 was indeed co-purified with the stalled MCP as revealed by immunoblots using a newly produced anti-TbTim15 antibody (*Fig. S2A*). Thus, TbTim15 is the only candidate, which is part of both, the active presequence^32^ and the active carrier translocase (*Fig. S1*).

Finally, we added Tb927.8.7010 to our candidate list, even though it was enriched only fourfold in the TbPam27-myc SILAC CoIP, because it was highly enriched in the CoIP of the of the active presequence translocase but not associated with the carrier translocase (*Fig. 1A, Fig. S1A*).

The domain structure of all six new putative TIM or PAM subunits were analysed *in silico* (*Fig. 1C*). TbTim15 contains four CX_9_C motifs (C=cysteine, X=any amino acid but cysteine), which are typically found in IMS proteins^40,42^. In the AlphaFold structure prediction of TbTim15 (*Fig. S2B*)^43^ each twin CX_9_C motif is positioned in a loop, which would allow for the formation of two intramolecular disulfide bridges. Such disulfide bridges are known to trap the protein in the IMS. In both, yeast and *T. brucei,* the import of proteins into the IMS depends on TbErv1^40^. Moreover, in *T. brucei,* the MICOS subunit TbMic20 has also been implicated in import of IMS proteins^44,45^. Interestingly, in previous quantitative proteomics studies, TbTim15 was shown to be downregulated 3.65-fold upon TbErv1 RNAi^40^ and 2.36-fold upon TbMic20 RNAi^45^. We confirmed these findings by following the steady state levels of TbTim15 in previously generated TbErv1^40^ and TbMic20^45^ RNAi cell lines over several days of induction (*Fig.2AB*). Thus, TbTim15 most likely is an IMS protein.

For TbTim20, none of various online prediction tools^46–50^ recovered any recognizable domains (*Fig. 1C*). 7010, 1010, 16750 and 3450, all contain a single predicted TMD^50^ and for 1010, an N-terminal mitochondrial targeting signal (MTS) was predicted^51^. In addition, 7010 and 1010 contain predicted J-domains at their N-termini (*Fig. 1C*)^46,47^.

Basic Local Alignment Tool (BLAST)^52^ and secondary structure similarity analysis by HHPred^49^ showed that all candidates are conserved among kinetoplastids. TbTim15, TbTim20, 16750, and 3450 show no obvious similarities to proteins in *S. cerevisiae* or humans. For 7010 and 1010 the similarities detected to proteins of other species were limited to the respective J-domains.

### All candidates are mitochondrial integral-membrane or partially membrane-associated proteins

All our candidate proteins are components of the previously defined mitochondrial importome^40^ and all, except 3450, are components of an earlier identified cluster of IM proteins^53^. To confirm the mitochondrial localization of the six candidates experimentally, cell lines allowing the inducible ectopic expression of C-terminally HA-tagged versions of the proteins were generated. These cell lines were subjected to digitonin extractions allowing the separation of a mitochondria-enriched pellet from a cytosolic fraction. As shown in *Fig. S3*, all HA-tagged candidates co-fractionate with the mitochondrial marker ATOM40 in such an experiment. These findings were confirmed by immunofluorescence microscopy analysis, in which the HA-tagged candidates co-localized with ATOM40 (*Fig. S4*). Moreover, alkaline carbonate extractions of the mitochondria-enriched pellets showed that, except for the IMS-localized TbTim15-HA, all candidates were recovered in the pellet together with ATOM40. This suggests they are integral membrane proteins or strongly membrane-associated proteins (*Fig. S3*). However, while TbTim15-HA was mainly found in the soluble fraction a small portion was also found in the pellet suggesting TbTim15 is at least partially associated with a mitochondrial membrane. The pellet portion of TbTim15 does not represent an insoluble protein aggregate as it could be solubilized in 1% Triton X-100 (*Fig. S3*). These results suggest that a fraction of the IMS-localized TbTim15 is tightly membrane-associated.

TbTim20 is part of the IM proteome^53^, but it has no predicted TMD (*Fig. 1C*). The fact that it was recovered in the pellet fraction of an alkaline carbonate extraction (*Fig. S3*) suggests that it is closely associated with the IM, despite the absence of a predicted TMD. Moreover, as in the case of TbTim15, the pellet can be solubilized with Triton X-100 indicating it is not an insoluble protein aggregate. Concerning 7010, 1010 and 16750, previous proteomic analyses^40,53^, the predicted TMDs (*Fig. 1C*) and the experimental data in *Fig. S3* and *Fig. S4* suggest that they all are integral IM proteins. 3450 is a mitochondrial membrane protein that was not detected in the abundance profiling that led to the IM proteome^53^.

### TbTim15 and TbTim20 are associated with the TIM complex

To confirm that our selected candidates are associated with the TIM complex and interact with TbPam27, the HA-tagged candidate proteins were used as baits in anti-HA CoIP experiments. The resulting immunoblots were probed with anti-HA antibodies and the previously established antisera against the TIM subunits TbTim17, TimRhom I, as well as the small Tim chaperone Tim9^32^. Furthermore, newly produced antibodies recognizing TbPam27 and TbTim15 were used (for validation of antibodies see *Fig. S2AC*). In line with their central position in the TIM complex protein interaction network (*Fig. 1B*), TbTim15-HA and TbTim20-HA CoIPs efficiently recovered TbPam27, as well as all tested TIM subunits (*Fig. 3A*). Thus, we conclude that TbTim15 and TbTim20 are associated with TbPam27 and the entire TIM com-plex.

**Figure 2.**
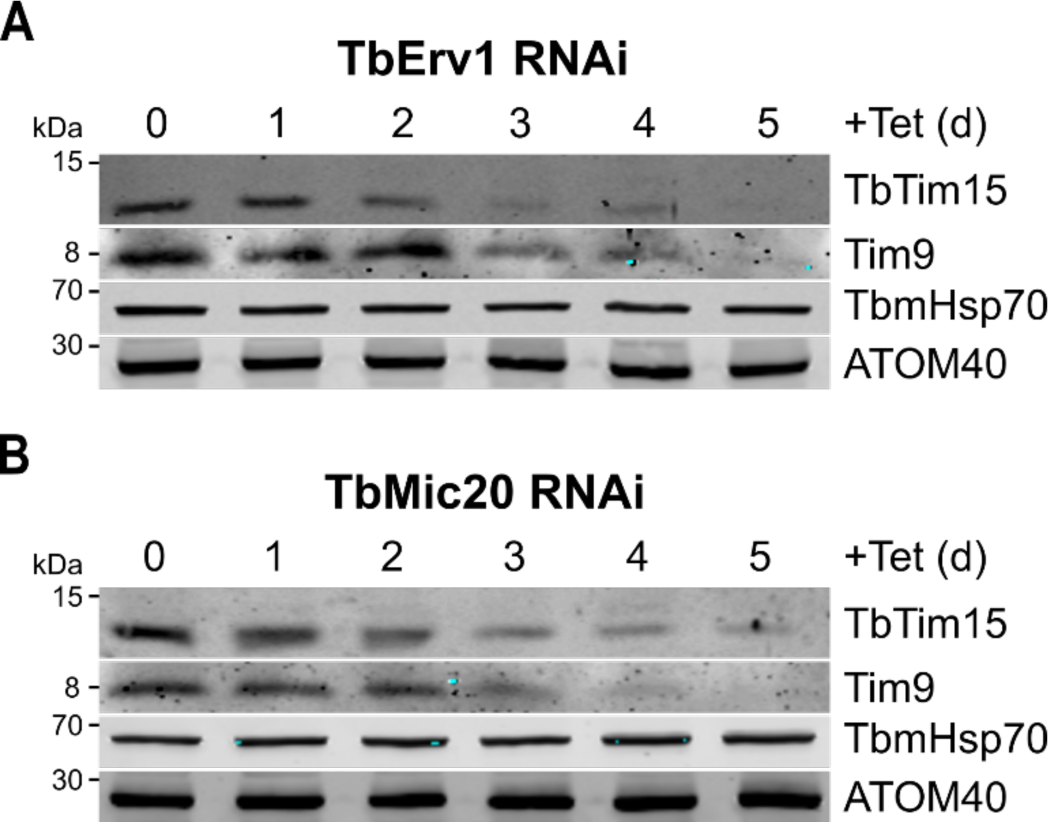
-TbTim15 is an intermembrane space protein. Immunoblot analysis of steady-state protein levels of TbTim15 in **(A)** TbErv1 and **(B)** TbMic20 RNAi background over five days of induction. Tim9, TbmHsp70 and ATOM40 were used as markers for intermembrane space, matrix, or outer membrane proteins, respectively.

**Figure 3.**
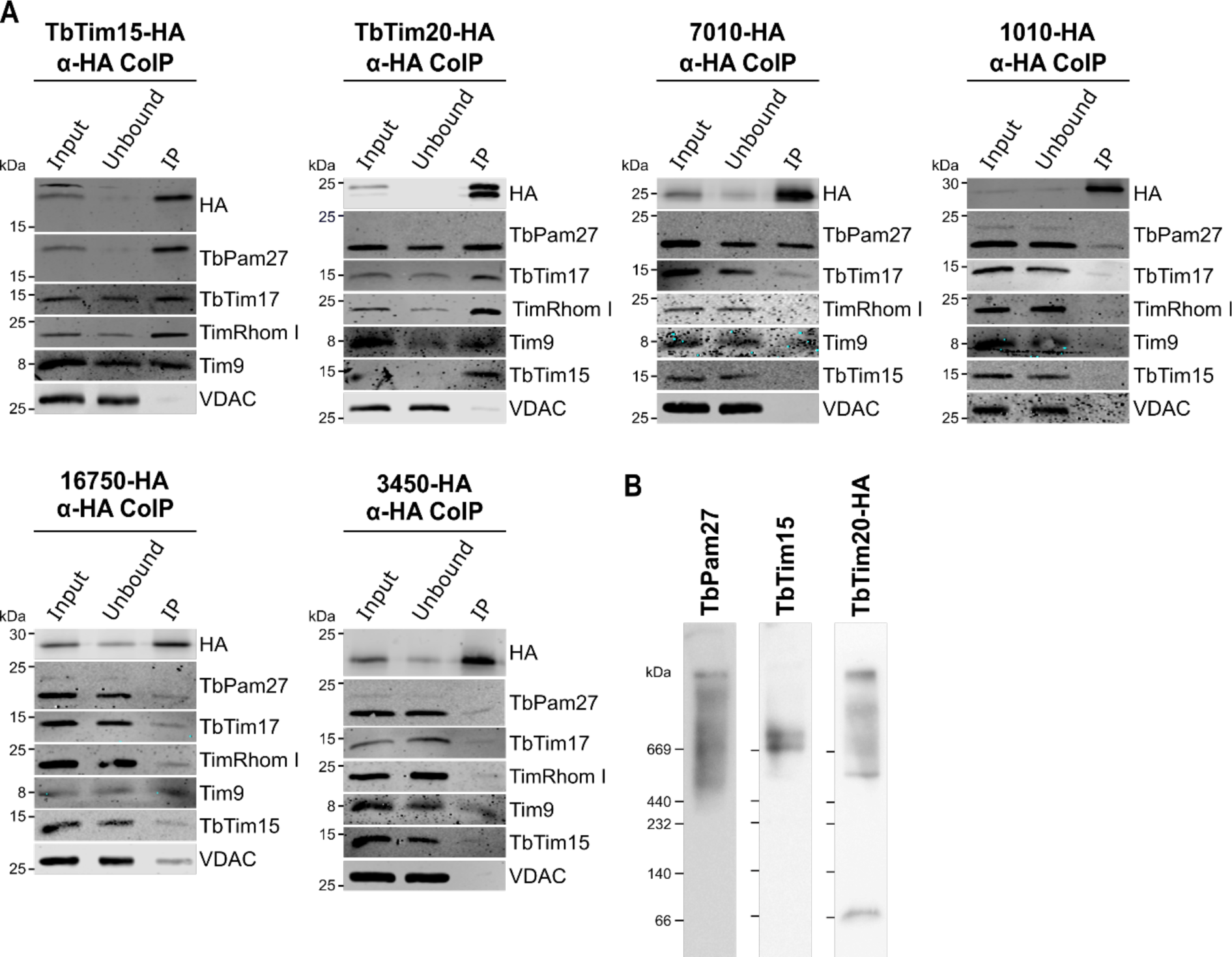
-TbTim15 and TbTim20 are associated with the TIM complex. **(A)** Cell lines expressing the C-terminally HA-tagged candidate proteins were subjected to co-immunoprecipitation (CoIP) experiments. 5% of mitochondria-enriched fractions (Input), 5% of unbound proteins (Unbound) and 100% of the final eluates (IP) were analysed by immunoblotting. The immunoblots were probed with anti-HA antibodies, antisera against TbPam27 and TbTim15, as well as the TIM subunits TbTim17, TimRhom I, the small Tim Tim9, and the voltage-dependent anion channel VDAC. **(B)** Blue native (BN)-PAGE analysis of TbPam27, TbTim15, and TbTim20-HA complexes in mitochondria-enriched fractions. Blots were probed with anti-TbPam27, anti-TbTim15, and anti-HA antibodies.

In the 7010-HA CoIP (*Fig. 3A*), TbPam27 was found to be enriched, as expected from the TbPam27-myc SILAC CoIP (*Fig. 1A, Table S1*) confirming the reciprocal interaction between the two proteins. However, it did not interact with any of the other tested TIM subunits, and thus, does not show up in the TIM complex protein interaction network (*Fig. 1B*). Therefore, 7010 seems to be only recruited to the TIM complex, when a presequence-containing substrate is being imported.

CoIP experiments in which 1010-HA, 16750-HA and 3450-HA were used as the baits, did not enrich for TbPam27 or any other of the tested TIM subunits (*Fig. 3A*). This is surprising because all of them were enriched more than fivefold in the TbPam27 SILAC CoIP (*Fig. 1A, Table S1*) and significantly enriched in more than one TIM subunit SILAC CoIP performed in previous studies (*Fig. 1B*). The reason for this is unclear but 16750 and 3450 localize to the periphery of the TIM complex protein interaction network indicating a weaker association with the TIM complex (*Fig. 1B*).

Because they are the only candidates clearly associated with TbPam27 and the TIM complex, we focused on TbTim15 and TbTim20 for further analysis. Blue native (BN)-PAGE, which allows the size separation of multiprotein complexes under native conditions, showed that TbTim15, TbTim20-HA and TbPam27 are all present in high molecular weight (HMW) complexes (*Fig. 3B, Fig. S2D*). Interestingly, while TbTim15 is present in two distinct complexes of approximately 700 kDa, the complexes formed by TbPam27 and TbTim20 overlap with the ones containing TbTim15, but are highly heterogenous.

In summary, these results show that TbTim15 and TbTim20 are present in the PAM/TIM/small Tim supercomplex. Thus, TbTim15 and TbTim20 are novel subunits of the trypanosomal IM protein import machinery.

### TbTim15 is essential for mitochondrial protein import

To learn more about the functions of TbTim15, TbTim20, 7010, 1010, 16750 and 3450, inducible RNAi cell lines were established. *Fig. 4A* shows that RNAi-induced knockdown of TbTim15 led to a strong growth retardation after two days of induction and a complete growth arrest at later timepoints. Knockdown of TbTim20, 7010 and 1010 does not affect normal growth. Thus, TbTim20, within the limits of RNAi analysis, is not essential for normal growth even though it is clearly associated with both TIM and PAM subunits (*Fig. 1AB, Fig 3*). Finally, RNAi-mediated ablation of 16750 and 3450 causes a slight reduction in growth rates between three and four days after RNAi induction (*Fig. 4A*).

**Figure 4.**
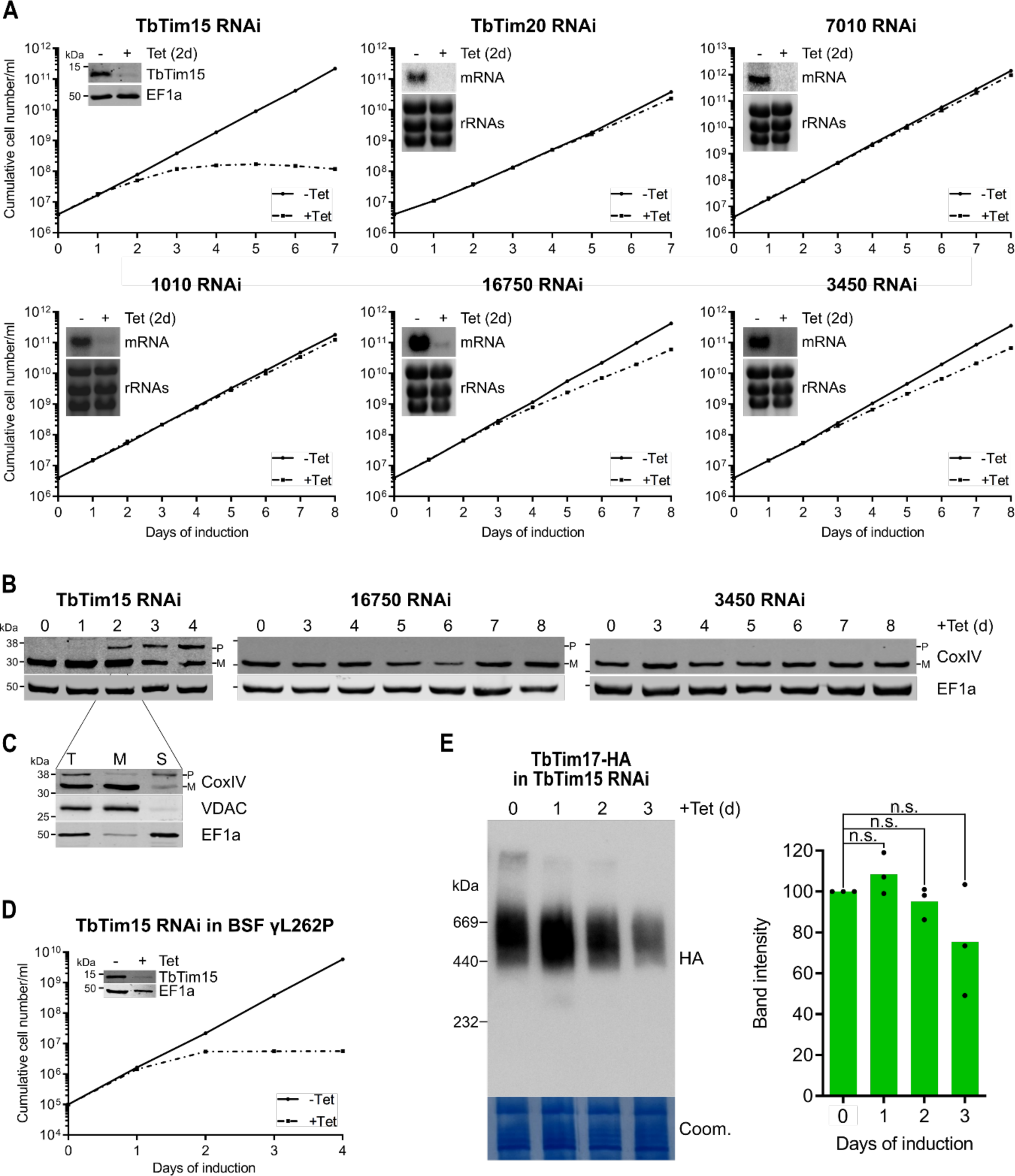
-TbTim15 is essential for mitochondrial protein import. **(A)** Growth curves of uninduced (-Tet) and induced (+Tet) RNAi cell lines. Inset in the growth curve of the TbTim15 RNAi cell line shows immunoblot of whole cell extracts of uninduced and two days induced RNAi cells. The blot was probed with anti-TbTim15 antibody and EF1a as loading control. Insets in remaining growth curves show north-ern blots of total RNA extracted from uninduced and two days induced cells, which were probed for the corresponding mRNAs. Ethidium bromide-stained ribosomal RNAs (rRNAs) serve as loading con-trols. **(B)** Immunoblot analysis of steady-state protein levels of cytochrome oxidase subunit 4 (CoxIV) in whole cell extracts of the indicated RNAi cell lines. CoxIV precursor (P) and mature form (M) are indicated. Elongation factor 1 alpha (EF1a) serves as loading control. **(C)** Immunoblot analysis of total TbTim15 RNAi cells (T), as well as digitonin-extracted, mitochondria-enriched (M), and soluble cytosolic (S) fractions. Blots were probed with anti-CoxIV antibodies and antisera against VDAC and EF1a, which serve as mitochondrial and cytosolic markers, respectively. **(D)** Growth curve of uninduced (-Tet) and induced (+Tet) bloodstream form (BSF) γL262 cell lines ablating TbTim15. Inset shows immunoblot of whole cell extracts of uninduced and two days induced RNAi cells. The blot was probed with anti-TbTim15 antibody and EF1a as loading control. **(E)** Left: BN-PAGE analysis of the TIM complex in cell lines expressing TbTim17-HA in the background of TbTim15 RNAi. The resultant immunoblot was probed with anti-HA antibody. Coomassie-stained gel section (Coom.) serves as loading control. Right: Densitometric quantification of the BN-PAGE. Green bars correspond to the mean of three independ-ent biological replicates. Levels in uninduced cells were set to 100%. n.s.: not significant, as calculated by an unpaired two-sided t-test.

Next, we tested whether TbTim15, 16750 and 3450, for which RNAi induction caused a change in the growth rate, are involved in mitochondrial protein import. To that end, we analysed steady-state levels of cytochrome c oxidase subunit 4 (CoxIV) in whole cell extracts of the respective RNAi cell lines (*Fig. 4B*). Accumulation of the unprocessed precursor of CoxIV in the cytosol, which still contains its N-terminal presequence, is a hallmark of a general mitochondrial protein import defect^31,32,39^. In the TbTim15 RNAi cell line, but not in the other two tested cell lines, we observed an accumulation of CoxIV precursor after two days of induction, which corresponds to the onset of the growth phenotype (*Fig. 4A*).

Digitonin fractionation showed that while the mature form of CoxIV almost exclusively co-fractionated with the mitochondrial marker, the CoxIV precursor was found in the cytosolic fraction (*Fig. 4C*), as would be expected in the case of an import defect. Moreover, immunofluorescence microscopy of ATOM40- and Mitotracker-stained TbTim15 RNAi cells confirmed that the general mitochondrial morphology remained unchanged and that the mitochondrial membrane potential was still intact at the timepoint the import phenotype becomes apparent (*Fig. S5*).

*T. brucei* has a complex life cycle and alternates between an insect vector, the Tsetse fly, and a mammalian host. In the procyclic form (PCF) in the fly, the mitochondrion is capable of oxidative phosphorylation (OXPHOS). In contrast, mitochondria of the bloodstream form (BSF) lack the respiratory complexes, except for the ATP-synthase, which functions in reverse^54^. However, both the PCF and the BSF, depend on mitochondrial protein import^55^. Results obtained with an inducible TbTim15 RNAi cell line of the BSF strain γL262P^56^ showed that TbTim15, as expected for a general import factor, is also essential for normal growth of BSF trypanosomes (*Fig. 4D*).

TbTim15 could be required either for the assembly of the TIM complex or directly mediate protein import. BN-PAGE analysis of a TbTim15 RNAi cell line expressing a C-terminally HA-tagged version of the TIM core subunit TbTim17 showed that the formation and stability of the TIM complex was not significantly affected for at least three days after RNAi induction (*Fig. 4E*). This suggests that TbTim15 does not mediated TIM complex assembly and instead has a more direct role in mitochondrial protein import.

### TbTim15 is required to stall the presequence and the carrier pathway intermediates

Using artificial import substrates that can be arrested in either the presequence or the MCP pathway, it could be shown that some TIM subunits are specifically associated with the active presequence translocase but absent from the MCP translocase form of the single trypanosomal TIM complex^32^. The presequence pathway intermediate is formed by the expression of a C-terminally tagged presequence-containing substrate that is fused to dihydrofolate reductase (DHFR). Addition of the folate-analogue aminopterin (AMT) prevents import of the C-terminal part of the chimeric protein as it tightly binds to the DHFR moiety. This results in a substrate that is stuck in the import channels across both the OM and the IM. The MCP pathway intermediate is formed by expression of truncated version of a tagged MCP that lacks the two N-terminal TMDs, which causes its accumulation in the carrier translocase. Formation of the two import intermediates can be monitored by BN-PAGE analysis.

To find out whether TbTim15 can be assigned to either the presequence or the carrier pathway, we expressed the two modified import substrates in the background of TbTim15 RNAi. *Fig. 5* shows that after two days of RNAi induction, at the onset of the growth phenotype (*Fig. 4A*) and at a time when the TIM complex was still intact (*Fig. 4E*), the formation of both substrate-containing translocase complexes was significantly reduced. These findings demonstrate that TbTim15 is required for the formation of both, the presequence and the carrier pathway intermediate. Hence, it is probably involved in both pathways. These results also suggest that TbTim15 is rather a component of the TIM complex than of the PAM module. Importantly, the same assay had previously been used to show that TbPam27, as expected for a PAM subunit, is selectively required for the formation of the presequence but not the carrier pathway intermediate^39^.

**Figure 5.**
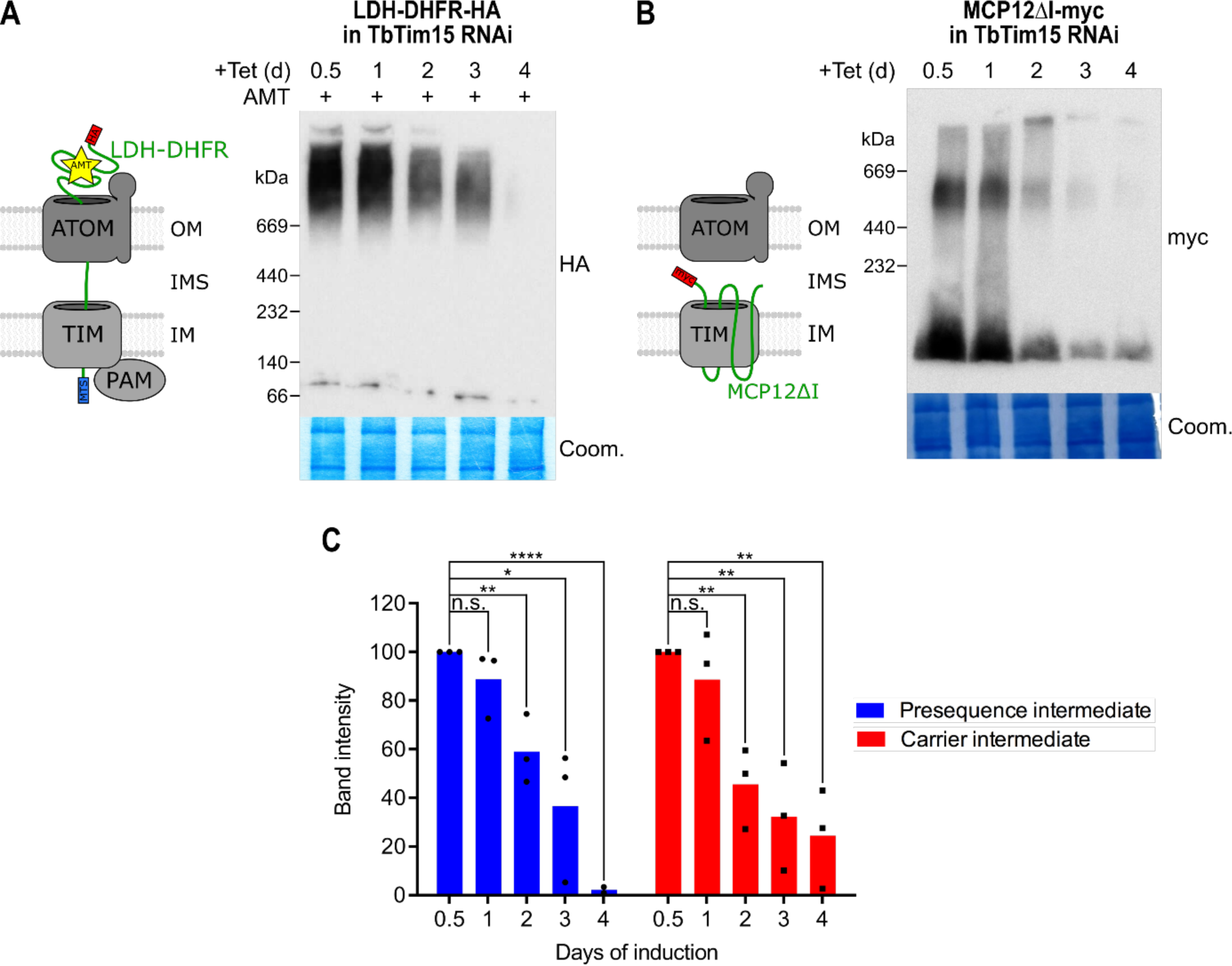
-TbTim15 is required to stall the presequence and the carrier pathway intermediates. **(A)** Left: Schematic representation of the stalled presequence pathway intermediate induced *in vivo* by expression of the LDH-DHFR-HA fusion protein in presence of aminopterin (AMT). Right: BN-PAGE analysis of the presequence pathway intermediate in mitochondria-enriched fractions of cells expressing LDH-DHFR-HA in the background of TbTim15 RNAi. Cells were induced (+Tet) for 0.5-4 days (d) and grown in presence of AMT for 0.5 days prior to the experiment. The resultant immunoblot was probed with anti-HA antibody. Coomassie-stained gel section (Coom.) serves as loading control. **(B)** Left: Schematic representation of the import-arrested carrier intermediate induced *in vivo* by the expression of the truncated myc-tagged mitochondrial carrier protein 12 (MCP12ΔI-myc). Right: BN-PAGE analysis of the carrier intermediate in mitochondria-enriched fractions of cells expressing MCP12ΔI-myc in the background of TbTim15 RNAi. Cells were induced (+Tet) for 0.5-4 days (d). The resultant immunoblot was probed with anti-myc antibody. Coom. serves as loading control. **(C)** Densitometric quantification of the BN-PAGE signals to compare the amount of stalled LDH-DHFR-HA (as shown in A) and MCP12ΔI-myc (as shown in B) in TbTim15 RNAi background in the respective high molecular weight complexes. The levels after 0.5 days of RNAi induction were set to 100%. Blue (presequence intermediate) and red (carrier intermediate) bars represent the mean of three independent biological replicates, respectively. n.s.: not significant, *: p-value<0.05, **: p-value<0.01, ****: p-value<0.0001, as calculated by an unpaired two-tailed t-test.

## Discussion

*T. brucei* contains a non-canonical single TIM complex that mediates the import of prese-quence-containing proteins and of MCPs^32^. To translocate presequence-containing proteins, the TIM complex associates with an unusual PAM that contains the kinetoplastid-specific TbPam27^39^. Here, we used TbPam27 as the bait in a SILAC CoIP experiment, which together with a re-analysis of previously published SILAC CoIPs of various TIM complex subunits, revealed six new candidates for TIM subunits or TIM-associated proteins (*Fig. 1AB*). <u>These can-didates are</u> all kinetoplastid-specific, mitochondrial proteins (*Fig. S3*)^40^. RNAi-mediated depletion of these proteins showed that only TbTim15 is essential for normal growth and that depletion of the other five candidates either did not or only marginally impede growth of PCF trypanosomes (*Fig. 4A*).

Thus, we focused our studies mainly on TbTim15. It contains twin CX_9_C motifs (*Fig. 1C, Fig. S2B*) that are characterized by two α-helices forming an antiparallel α-hairpin that is covalently paired by two disulfide bridges^57^. Twin CX_9_C motif-containing proteins generally localize to the IMS. This is also the case for TbTim15, which requires the IMS import factors TbErv1^40^ and TbMic20^45^ for its import (*Fig. 2*). Twin CX_9_C motif-containing proteins perform diverse mitochondrial functions. Many are involved in OXPHOS as non-catalytic subunits or assembly factors of complex I and complex IV of the respiratory chain. Others regulate mitochondrial morphology as MICOS subunits or are involved in maintaining lipid homeostasis^57^. Whereas most twin CX_9_C motif-containing proteins have a single twin CX_9_C motif, TbTim15 contains two. Other examples for proteins with quadruple CX_9_C motifs include the mammalian complex I subunit NDUFA8^58^ and the mammalian MICOS subunit Mic14^59^.

Three lines of experimental evidence indicate that the trypanosomal TbTim15 functions in mitochondrial protein import: i) While not having an obvious predicted TMD (*Fig. S3*), TbTim15 is constitutively associated with the single PAM/TIM/small TIM supercomplex (*Fig. 1AB, Fig. 3, Fig. S1*), regardless of whether it is engaged in import of presequence-containing proteins or MCPs (*Fig. S1*); ii) depletion of TbTim15 results in the accumulation of uncleaved precursor of CoxIV (*Fig. 4BC*), and prevents the formation of import-arrested substrates in both, the presequence as well as the carrier translocases (*Fig. 5*); and iii) TbTim15 is essential for normal growth of both, the PCF and the BSF of trypanosomes (*Fig. 4AD*). BSF parasites, except for the ATP synthase, lack the respiratory complexes, and therefore, are not capable of OXPHOS^60^. This indicates that the role of TbTim15, unlike many twin CX_9_C motif-containing proteins in other organisms, cannot be restricted to OXPHOS. Rather it must be involved in a more general process that is essential in both life cycle stages, such as mitochondrial protein import.

Presently, the precise function of TbTim15 in mitochondrial protein import is unknown. There are many possibilities, including that it may function as an IMS-localized chaperone, such as the six previously identified small TIM proteins^37,61^, or as a receptor for substrates of the TIM complex. Intriguingly, there is a well-studied twin CX_9_C motif-containing protein, termed Mia40 that is involved in mitochondrial protein import in yeast and mammals. Mia40 together with Erv1 forms a disulfide relay, which imports small cysteine motif-containing proteins into the IMS. In this relay, Mia40 performs two distinct functions: it serves as a receptor for its substrates, and it oxidizes their cysteines^62^. The electrons liberated in the oxidation are subsequently transferred to cytochrome c by the sulfhydryl oxidase Erv1. While Erv1 and Mia40 are conserved in most eukaryotes, *T. brucei* and its relatives lack a Mia40 orthologue^63^. It is therefore unclear whether the trypanosomal Erv1 directly oxidizes IMS proteins or, as has been suggested in plants^64^, whether Mia40 has been replaced by another protein^65,66^.

It would be interesting to investigate whether TbTim15 could, at least partially, take over the function of the lacking Mia40. In yeast, the essential receptor function of Mia40 is mediated by the pocket formed by the twin CX_9_C motif, whereas the oxidase activity requires the CPC motif in the N-terminal domain of the protein^62^. TbTim15 lacks a CPC motif, and therefore, likely has no oxidase activity. Yet, it does have two twin CX_9_C motifs (*Fig. 1C, Fig. S2B*), which could form pockets recognizing imported proteins. However, in contrast to yeast Mia40, TbTim15 is associated with the TIM complex (*Fig. 1AB, Fig. 3*) and involved in both, the presequence and the MCP import pathway (*Fig. S1, Fig. 5*). This suggests that, should TbTim15 indeed be an import receptor, it would recognize different or a wider range of substrates than Mia40, including presequence-containing as well as MCPs. Intriguingly, it has recently been proposed that also yeast Mia40 might be involved in import of non-typical substrates^67–69^. Moreover, trypanosomes lack a Tim50 orthologue, which serves as an import receptor for the TIM23 complex of yeast and mammals, and no other receptor subunit has been identified^32^. Contrary to depletion of any of the integral membrane TIM subunits^32^, lack of TbTim15 seems to not directly interfere with the assembly or the stability of the TIM complex (*Fig. 4E*). This is reminiscent of the two receptors ATOM46 and ATOM69 of the ATOM complex of trypanosomes^31^, and of the Tom20 and Tom70 receptors of the yeast and mammalian TOM complexes^70^, whose absence does not affect the integrity of the (A)TOM complexes.

It has recently been suggested that the IMS-localized trypanosomal protein TbTim54, as TbTim15, is associated with the trypanosomal TIM complex (note that despite its name, TbTim54 is not related to the yeast TIM22 complex subunit Tim54)^34,71^. In contrast to TbTim15, it was proposed that TbTim54 mediates the assembly of the TIM complex and the import of at least a subset of MCPs^71^. However, unlike expected for a TIM complex assembly factor, knockdown of TbTim54 only marginally affects cell growth^34^. Moreover, TbTim54 was neither enriched in any of the SILAC pulldowns of the core TIM subunits (TbTim17, TbTim42, ACAD)^32,39^, the small TIM protein TbTim13^32^, nor the PAM subunit TbPam27 tested in the present study (*Table S1*). Thus, the question how closely TbTim54 is associated with the trypanosomal TIM complex remains open at present.

In summary, we show that TbTim15 is a new kinetoplastid-specific subunit of the single trypanosomal TIM complex that is involved in both the presequence as well as the mitochondrial carrier import pathway. TbTim15 is essential in the PCF and the BSF of trypanosomes. Our results are consistent with the idea that TbTim15 might be an IMS-localized protein import receptor. However, direct demonstration of the receptor function of TbTim15 and the identification of which substrates it recognizes require extensive biochemical analyses, which are beyond the scope of the present study.

## Material and methods

### Transgenic cell lines

Transgenic *T. brucei* cell lines are based on the procyclic form (PCF) strain 29-13^72^ or the BSF strain F1γL262P^56^. PCF cells were grown in SDM-79^73^ at 27°C, BSF parasites were grown in HMI9^74^ at 37°C, both supplemented with 10% (v/v) foetal calf serum (FCS).

To produce plasmids for ectopic expression of C-terminal triple HA-tagged TbTim15 (Tb927.2.4445), TbTim20 (Tb927.11.1620), Tb927.8.7010, Tb927.11.1010, Tb927.11.16750 and Tb927.8.3450 the complete ORF of the respective gene was amplified by PCR. The PCR product was subsequently cloned into a modified pLew100 vector^72,75^, which contains a puromycin resistance gene and a triple HA-tag^76^. The triple HA-tagged LDH-DHFR fusion protein and the triple c-myc-tagged truncated (nt 274-912) ORF of MCP12 (Tb927.10.12840) have been described previously^32^. Cell lines expressing C-terminally myc-tagged TbPam27 and HA-tagged TbTim17 have been described before as well^32,39^.

RNAi cell lines were generated using the same pLew100-derived vector described above and as described previously^39,77^. The RNAi targets the indicated nt of the ORF of TbTim15 (nt 4-394), TbTim20 (nt 58-492), 7010 (nt 65-464) and 3450 (nt 22-412) or the 3’ untranslated region (UTR) of 1010 (nt +4 -+292) and 16750 (nt +19 -+260). The TbErv1 RNAi^40^, TbMic20 RNAi^45^ and TbPam27^39^ RNAi cell lines have been described previously.

### Antibodies

Polyclonal rabbit antiserum against TbPam27 and TbTim15 was commercially produced (Eurogentec, Belgium) using aa 128-142 (RFTTRQHKSRISYDE) and aa 119-133 (VGLIQRQRGRQEQRR) as antigens, respectively. For immunoblots (WB) the TbPam27 antiserum was used at a 1:200 and the TbTim15 antiserum at a 1:250-500 dilution. Commercially available antibodies were: Mouse anti-c-myc (Invitrogen, dilution WB: 1:2’000) mouse anti-HA (Sigma-Aldrich, dilution WB: 1:5’000, dilution immunofluorescence (IF): 1:1’000) and mouse anti-EF1a (Merck Millipore, dilution WB 1:10’000). Antibodies previously produced in our laboratory are: polyclonal rabbit anti-ATOM40 (dilution WB: 1:10’000, dilution IF: 1:1’000), polyclonal anti-VDAC (dilution WB: 1:1’000), polyclonal rabbit anti-CoxIV (dilution WB: 1:1’000), polyclonal rabbit anti-Cyt C (dilution WB 1:100), polyclonal rabbit anti-Tim9 (dilution WB: 1:40), polyclonal rabbit anti-TimRhom I (dilution WB: 1:150) and polyclonal rat anti-TbTim17 (dilution WB: 1:300)^31,32,53^. Monoclonal anti-TbmHsp70 (dilution WB: 1:1’000) has been generated and described previously^78^. Secondary antibodies used: goat anti-mouse IRDye 680LT conjugated (LI-COR Biosciences, dilution WB: 1:20’000), goat anti-mouse Alexa Fluor 596 (ThermoFisher Scientific, dilution IF: 1’000), goat anti-rabbit IRDye 800CW conjugated (LI-COR Biosciences, dilution WB 1:20’000), goat anti-rabbit Alexa Fluor 488 (Invitrogen, IF: 1:1’000) and goat ani-rat IRDye 680LT conjugated (LI-COR biosciences, dilution WB 1:10’000). Immunoblots of BN-PAGE analysis were decorated with HRP-coupled goat anti-mouse or HRP-coupled goat anti-rabbit (both Sigma) as secondary antibodies (dilution: 1:5’000).

### Digitonin, alkaline carbonate and Triton X-100 extraction

Digitonin, alkaline carbonate and Triton X-100 extractions have been done as described before^77^.

### Co-immunoprecipitation (CoIP)

A mitochondria-enriched digitonin pellet from 1 x 10^8^ cells expressing the protein of interest was solubilized in a buffer containing 20 mM Tris-HCl (pH 7.4), 0.1 mM EDTA, 100 mM NaCl, 10% glycerol, 1X Protease Inhibitor mix (Roche, EDTA-free), and 1% (w/v) digitonin for 15 min at 4°C. After centrifugation (20’000 g, 15 min, 4°C) the lysate was transferred to 50 μl HA bead slurry (anti-HA affinity matrix, Roche) or 30 μl c-myc bead slurry (EZview red anti-c-myc affinity gel, Sigma), which had been equilibrated in wash buffer (20 mM Tris-HCl (pH 7.4), 0.1 mM EDTA, 1 mM NaCl, 10% glycerol, 0.2% (w/v) digitonin). Subsequent to incubation in an end-over-end shaker for at least one hour at 4°C, the supernatant containing the unbound proteins was removed. After washing the bead slurry three times with wash buffer, the bound proteins were eluted by boiling the resin for 5 min in 2% SDS in 60 mM Tris-HCl (pH 6.8). 5% of both the input and the unbound proteins, and 100% of the IP sample were analysed by SDS-PAGE and immunoblotting.

### Blue native (BN)-PAGE

BN-PAGE has been done as described before^31,32,39,77^.

For analysis of the stalled presequence pathway intermediate, LDH-DHFR expression was induced by tetracycline and cell cultures were supplemented with 1 mM sulfanilamide and 50 μM aminopterine (AMT), 0.5 days before the experiment^32,39^.

### Immunofluorescence (IF) microscopy

For analysis of the mitochondrial membrane potential, uninduced and induced TbTim15 RNAi cells were grown in presence of 500 nM MitoTracker Red CMXRos. As a negative control, un-induced cells were treated with 40 μM carbonyl cyanide m-chlorophenylhydrazone (CCCP) before incubation with MitoTracker.

For antibody-staining, cells were harvested, fixed with 4% paraformaldehyde in PBS and permeabilized with 0.2% Triton X-100 in PBS. Subsequently, the samples were decorated with primary antibodies for 1 hr. Washing with PBS was followed by incubation with secondary antibody for 1 hr. The cells were then postfixed in cold methanol and mounted using Vec-taShield containing 4ʹ,6-diamidino-2-phenylindole (DAPI) (Vector Laboratories). Images were acquired by a DMI6000B microscope and a DFC360 FX monochrome camera (both Leica Microsystems).

### RNA extraction and northern blotting

Acid guanidinium thiocyanate-phenol-chloroform extraction to isolate total RNA from uninduced and induced (two days) RNAi cells was done as described elsewhere^79^. To determine RNAi efficiency, the resulting RNA was used for RT-PCR or northern blotting as described previously^77^.

### SILAC CoIP experiments

Cells inducibly expressing TbPam27-myc were washed in PBS and resuspended in SDM-80^80^ supplemented with 5.55 mM glucose, 10% dialyzed FCS (BioConcept, Switzerland) and either light (^12^C_6_/^14^N_χ_) or heavy (^13^C_6_/^15^N_χ_) isotopes of arginine (1.1 mM) and lysine (0.4 mM) (Eurisotope). To make sure all proteins were completely labelled with heavy amino acids, the cells were grown in SILAC medium for six to ten doubling times. TbPam27-myc expressing cells were induced for 1 day. About 4 × 10^8^ uninduced and 4 × 10^8^ induced cells were harvested and mixed. Subsequently the mixture was extracted with Digitonin and subjected the co-IP protocol as describe above. The SILAC experiment was executed in three biological replicates including a label-switch and analysed by liquid chromatography-mass spectrometry (LC-MS).

### LC-MS and data analysis

Eluates of TbPam27-myc CoIPs were subjected to SDS-PAGE, followed by staining of the proteins using colloidal Coomassie Blue. Gel lanes were cut into 10 equal slices followed by reduction, alkylation, and tryptic in-gel digestion of proteins as described before^40^. LC-MS analyses were carried out using an UltiMate 3000 RSLCnano HPLC system (Thermo Scientific, Dreieich, Germany) connected to an Orbitrap Elite mass spectrometer (Thermo Scientific, Bremen, Germany). The RSLC system was operated with nanoEase M/Z Symmetry C18 pre-columns (Waters, Eschborn, Germany; length of 20 mm, inner diameter of 180 µm, flow rate of 5 – 10 µl) for washing and preconcentration of peptides and a nanoEase M/Z HSS C18 T3 Col analytical column (Waters; length of 250 mm, inner diameter of 75 μm, particle size of 1.8 μm, packing density of 100 Å, flow rate of 300 nl/min) for peptide separation. Peptides were eluted using a binary solvent system consisting of 4% DMSO/0.1% formic acid (FA) (solvent A) and 48% methanol/30% acetonitrile/4% DMSO/0.1% FA (solvent B) and a gradient ranging from 1 −65% B in 30 min, 65 −80% B in 5 min, and 3 min at 80% B. MS/MS data were obtained in data-dependent mode using the following parameters: full MS scans were acquired at a mass range of *m/z* 370 – 1,700 at a resolution of 120,000 (at *m/z* 400), a maximum automatic gain control (AGC) of 1 × 10^6^ ions, and a maximum injection time (IT) of 200 ms. A TOP15 method was applied for further fragmentation of multiply charged precursor ions by collision-induced dissociation. MS/MS scans were acquired with a normalized collision energy of 35%, an activation q of 0.25, an activation time of 10 ms, an AGC of 5,000, and a maximum IT of 150 ms. The dynamic exclusion time was set to 45 s.

Mass spectrometric raw data were processed using MaxQuant/Andromeda (^81,82^, version 1.5.5.1) and searched against all entries in the TriTryp database (https://tritrypdb.org, ver-sion 8.1) for peptide and protein identification using MaxQuant default parameters and Arg10 and Lys8 as heavy labels. The options “re-quantify” and “match between runs” were enabled. Protein identification was based on ≥ one unique peptide and a false discovery rate of 1% at peptide and protein level. SILAC-based relative protein quantification was based on unique peptides and ≥ one ratio count (i.e., SILAC peptide pair). Proteins enriched in TbPam27-myc complexes were identified by applying the rank sum method^83^, carried out using the package ‘RankProd’ (version 3.24.0;^84^) in R (version 4.2.2). *P* values and the percentage of false positives (i.e., q-values, referred to as adjusted *P* values) were determined for the enrichment of proteins quantified in at least two replicates. For a list of proteins identified and quantified, see *Table S1*.

### Data availability

The mass spectrometric data have been deposited to the ProteomeXchange Consortium^85^ (REF) via the PRIDE^86^ partner repository and are accessible using the dataset identifier PXD048257.

## Acknowledgements

We thank Julian Bender for assistance in bioinformatics data analysis. Work in the lab of A.S. was supported in part by NCCR RNA & Disease, a National Centre of Competence in Research (grant number 205601) and by project grant SNF 205200 both funded by the Swiss National Science Foundation.

## Supplementary Figures

**Figure S1.**
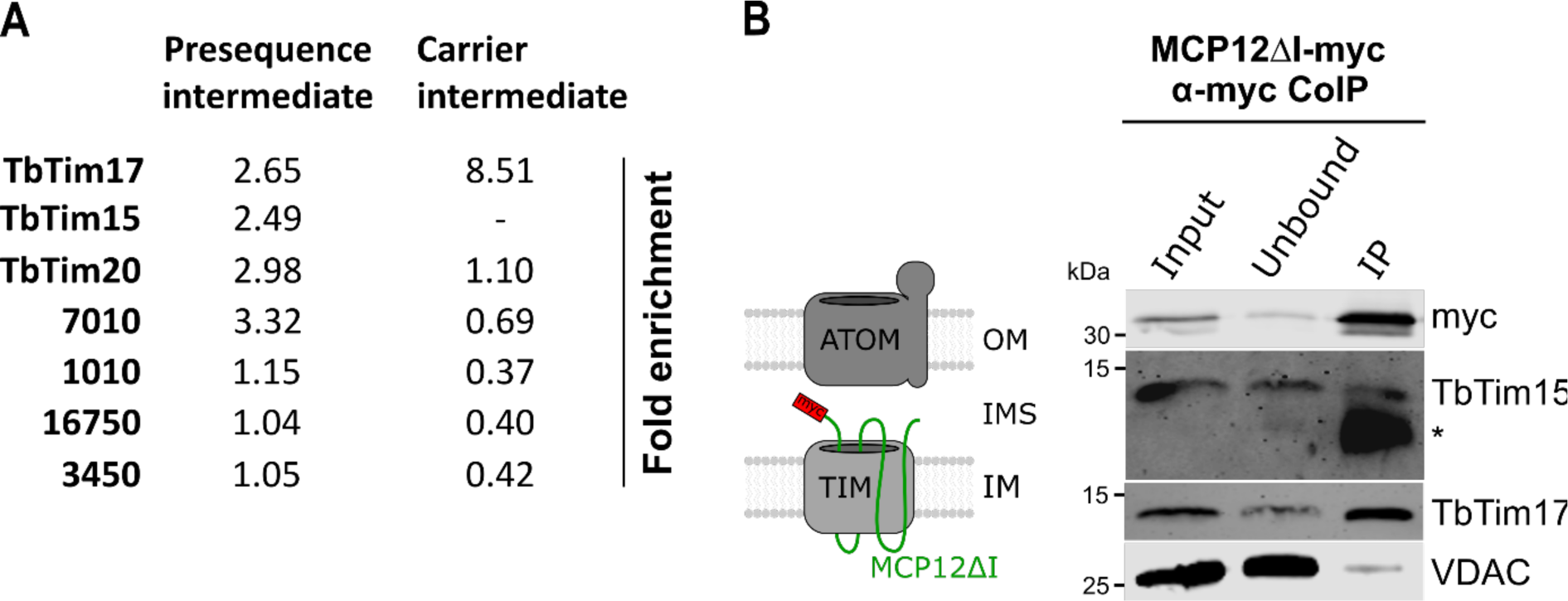
-Enrichment of TIM subunit candidates in stalled import intermediates. **(A)** Listing of enrichment factors (*P* value < 0.05) of candidate proteins in previously published SILAC CoIP experiments, in which stalled import substrates were used as baits. Enrichment factors of the TIM core subunit TbTim17 are listed for comparison. TbTim15 has not been detected in the carrier intermediate SILAC CoIP. **(B)** Left: Schematic representation of the stalled carrier intermediate formed after the expression of a truncated mitochondrial carrier protein (MCPΔI). Right: Digitonin-solubilized, mitochondria-enriched fractions of cells expressing MCP12ΔI-myc, were subjected to an anti-myc CoIP. 5% each of the crude mitochondrial fraction (Input) and unbound proteins (Unbound) and 100% of the final eluate (IP) were analysed by SDS-PAGE and immunoblotting. The resulting immunoblot was probed with anti-myc antibodies and antisera against TbTim15, TbTim17 and VDAC. *, unspecific band.

**Figure S2.**
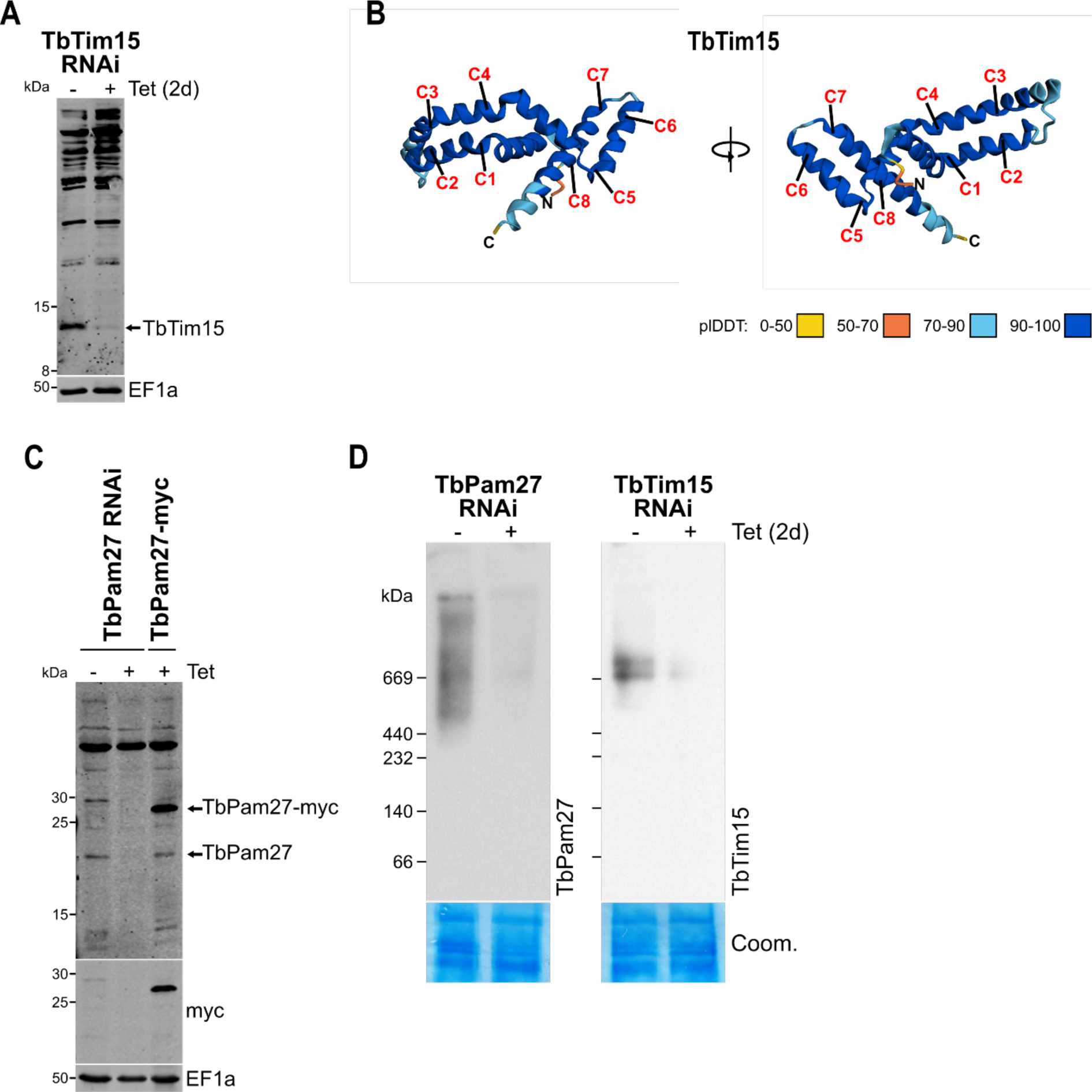
-Verification of antibodies and AlphaFold prediction. **(A)** SDS-PAGE and immunoblot analysis of whole cell extracts of uninduced and two days induced TbTim15 RNAi cells. Immunoblot was probed with anti-TbTim15 antibody. EF1a serves as loading control. **(B)** AlphaFold structure prediction of TbTim15. Cysteine residues (C) that are part of Cx_9_C motifs are indicated. **(C)** SDS-PAGE and immunoblot analysis of whole cell extracts of uninduced and two days induced TbPam27 RNAi cells (left two lanes) and of a cell line expressing myc-tagged TbPam27 (right lane). Immunoblot was probed with anti-TbPam27 antibody. EF1a serves as loading control. **(D)** BN-PAGE and immunoblot analysis of dig-itonin-extracted, mitochondria-enriched fractions of TbPam27 and TbTim15 RNAi cells. Immunoblots were probed with anti-TbPam27 and anti-TbTim15 antibody, respectively. Coomassie-stained gel sections (Coom.) serve as loading controls. Sections of the same blots and gels are shown in Fig. 3.

**Figure S3.**
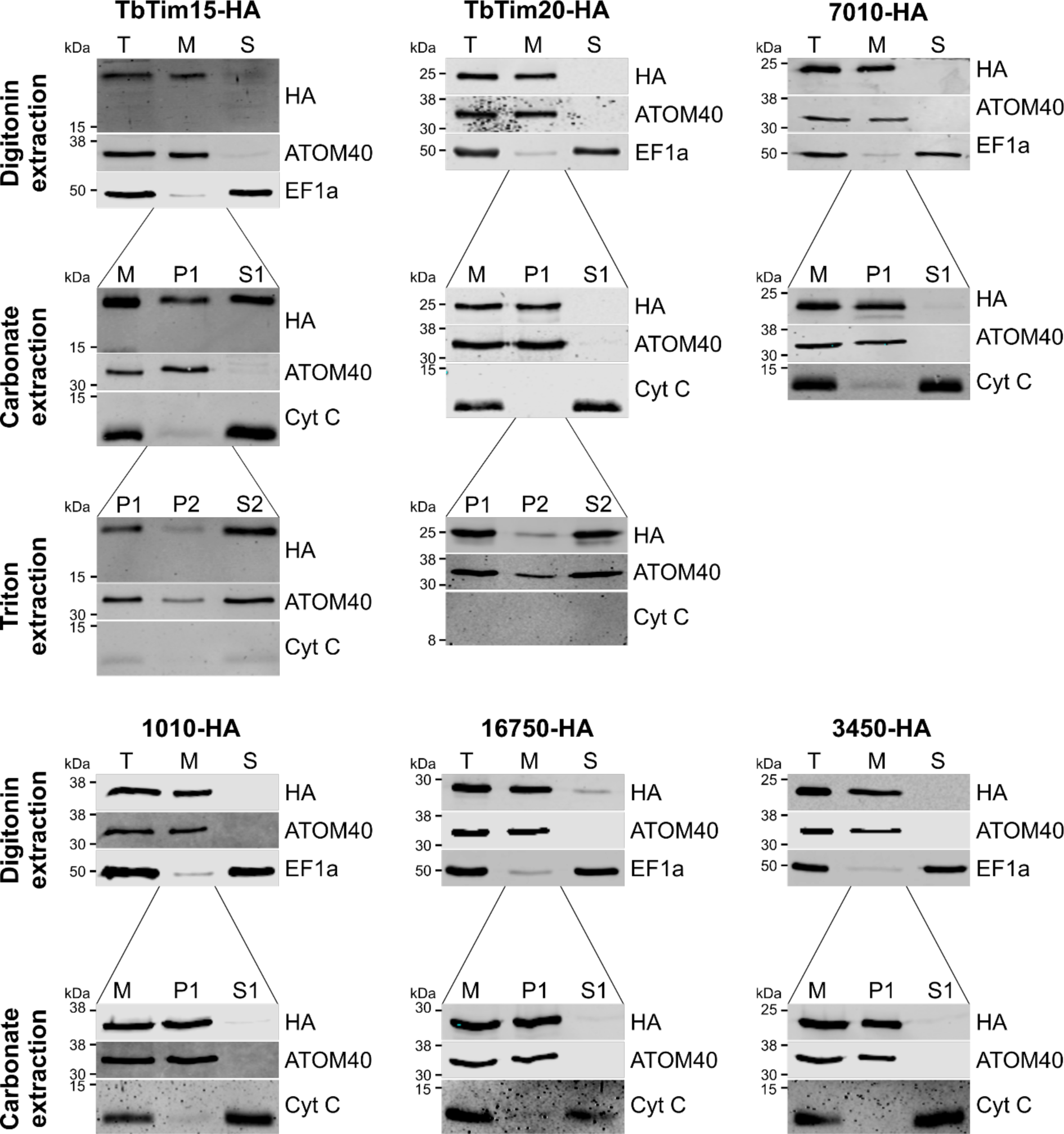
-TbTim15 is a partially soluble and partially membrane-associated, mitochondrial protein. Upper panels of each candidate: Immunoblot analysis of total cells (T), digitonin-extracted mitochon-dria-enriched (M), and soluble cytosolic (S) fractions of C-terminally HA-tagged candidate proteins. Blots were probed with anti-HA antibodies and antisera against ATOM40 and EF1a, which serve as mitochondrial and cytosolic markers, respectively. Second panels of each candidate: Digitonin-extracted, crude mitochondrial fractions (M) were subjected to an alkaline carbonate extraction resulting in a pellet enriched in membrane proteins (P1) and a supernatant fraction containing soluble proteins (S1). Immunoblots were probed with anti-HA and antisera against ATOM40 and cytochrome C (Cyt C), which serve as markers for integral membrane and soluble proteins, respectively. For TbTim15 and TbTim20, Triton X-100 extractions were done in addition (lowest panel): Half of the P1 fraction from the alkaline carbonate extraction was solubilized in 1% Triton X-100. Differential centrifugation resulted in a pellet (P2) and a soluble (S2) fraction. The immunoblots were probed with anti-HA, anti-ATOM40 and anti Cyt C antibodies.

**Figure S4.**
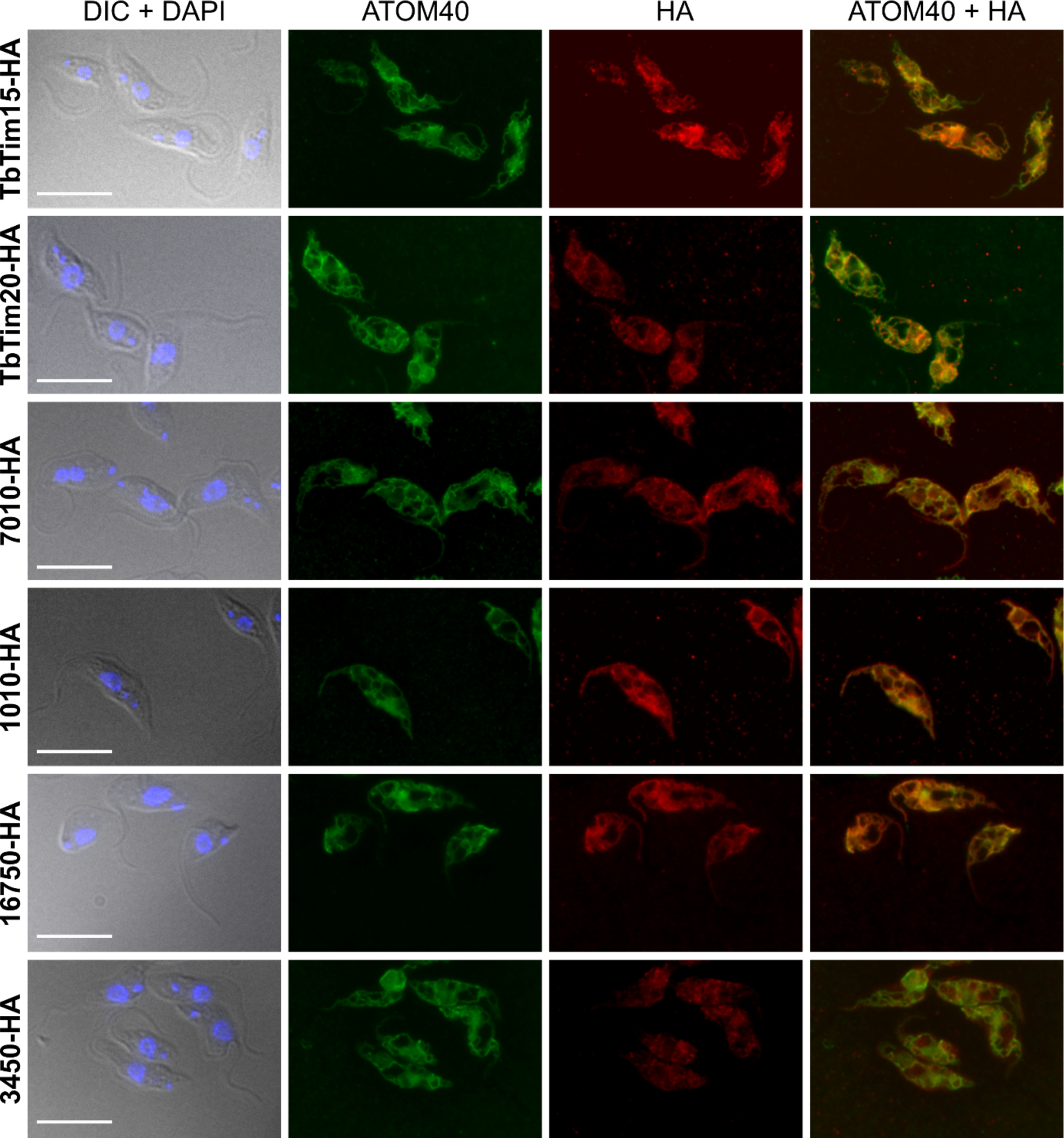
-TIM subunit candidates are all mitochondrial proteins. Differential contrast (DIC) and DAPI staining with corresponding ATOM40 and HA stainings of cell lines expressing the HA-tagged candidate proteins. ATOM40 serves as a mitochondrial marker. Bar; 10 μm.

**Figure S5.**
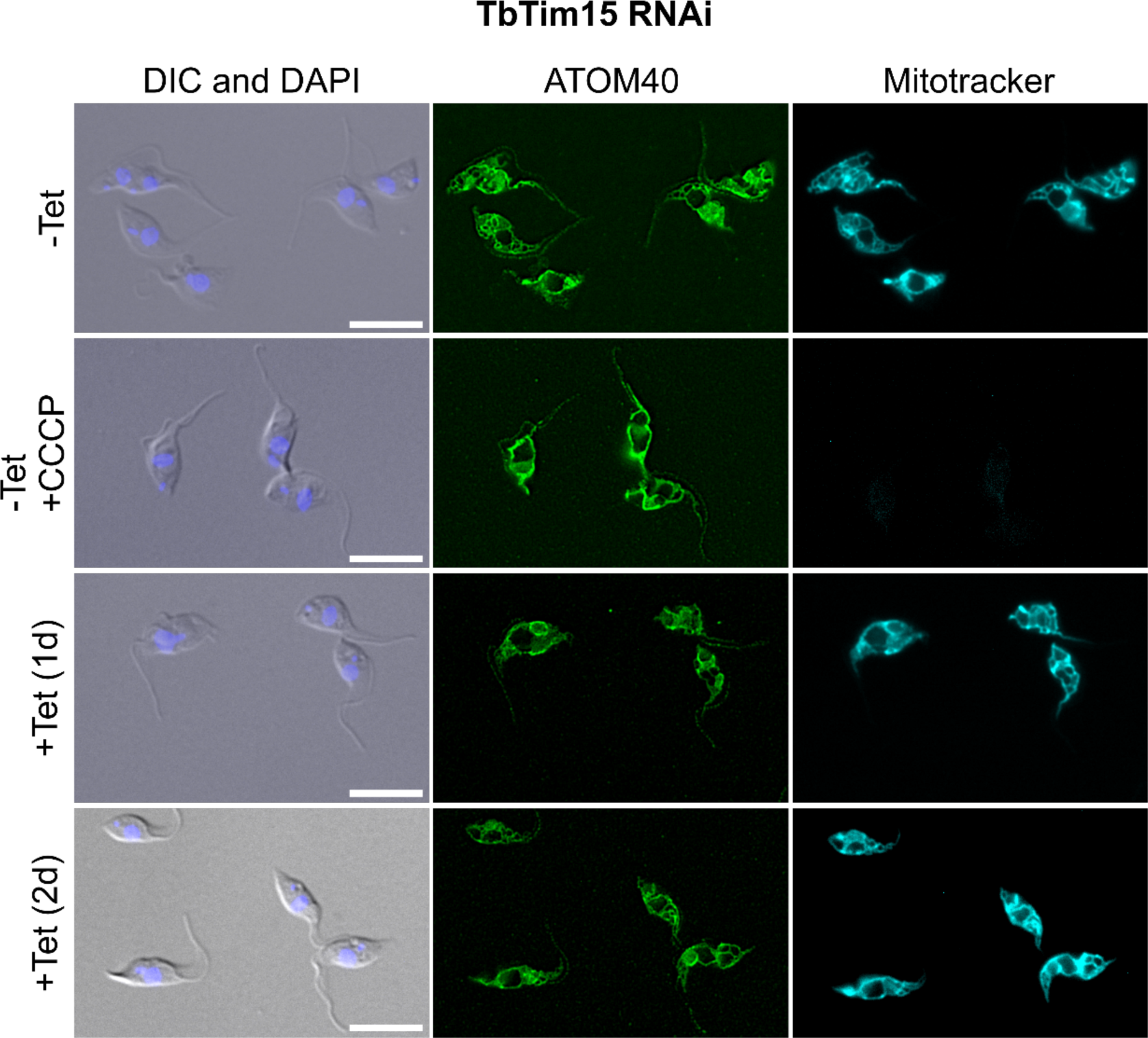
-TbTim15 RNAi does not abolish mitochondrial integrity or the mitochondrial membrane potential at early time points of induction. Differential contrast (DIC) and DAPI staining with corresponding ATOM40, as well as Mitotracker staining of uninduced (-Tet) and one and two days induced (+Tet) TbTim15 RNAi cells. Uninduced cells treated with 40 μM of carbonyl cyanide m-chlorophenylhy-drazone (CCCP) serve as control for cells that have lost the mitochondrial membrane potential. Bar; 10 μm.

